# raxmlGUI 2.0: a graphical interface and toolkit for phylogenetic analyses using RAxML

**DOI:** 10.1101/800912

**Authors:** Daniel Edler, Johannes Klein, Alexandre Antonelli, Daniele Silvestro

## Abstract

RaxmlGUI is a graphical user interface to RAxML, one of the most popular and widely used software for phylogenetic inference using maximum likelihood. Here we present raxmlGUI 2.0, a complete rewrite of the GUI which seamlessly integrates RAxML binaries for all major operating systems with an intuitive graphical front-end to set up and run phylogenetic analyses. Our program offers automated pipelines for analyses that require multiple successive calls of RAxML and built-in functions to concatenate alignment files while automatically specifying the appropriate partition settings. In addition to RAxML 8.x, raxmlGUI 2.0 also supports the new RAxML Next Generation. RaxmlGUI facilitates phylogenetic analyses by coupling an intuitive interface with the unmatched performance of RAxML.

## Introduction

Phylogenetic inference is a keystone in evolutionary biology research, providing the foundations for tackling a wide range of questions, from population dynamics to taxonomy of higher taxa (Felsenstein, 2003). RAxML is one of the most widely used programs in phylogenetic analysis, implementing extremely fast algorithms to analyze large datasets using maximum likelihood (Stamatakis, 2014). Despite the undisputed efficiency of RAxML, the program is only available through a command-line interface. This requires users to be familiar with the shell environment and to navigate through the ever-growing number of commands implemented in the program, which may exclude many potential users without such experience. RaxmlGUI (Silvestro and Michalak, 2012) is a graphical interface intended to facilitate phylogenetic analyses using RAxML by providing a graphical front-end to help users set up their analysis. Although this interface has been widely used, there are many areas of improvement in terms of accessibility, usage and performance.

Here, we present raxmlGUI 2.0, a complete rewrite of the raxmlGUI program. This version brings a new cross-platform design, novel functionalities and a seamless integration with both RAxML 8.2 and RAxML-NG, the new RAxML Next Generation (Kozlov et al., 2019). Similarly to its predecessor, raxmlGUI 2.0 is designed to be easy to use as a cross-platform stand-alone program that does not require the installation of additional software. It provides the user with an intuitive interface with access to the model settings required to setup and run a phylogenetic analysis, run on datasets comprising up to many thousand terminals (Lemoine et al., 2018). The GUI additionally provides a number of automated options to parse, concatenate, and partition alignments and to run analytical pipelines combining multiple RAxML calls. RaxmlGUI 2.0 targets a wide userbase ranging from beginners to advanced phylogeneticists seeking an easy access to the state-of-the-art analytical tools implemented in RAxML and RAxML-NG.

## Methods

The program comes with pre-compiled integrated versions of RAxML for the major operating systems (MacOS, Windows, Linux), including the PTHREADS and SSE3 versions (Stamatakis, 2014) allowing the user to run faster analyses using parallel computing, when multiple CPUs are available. Pre-compiled versions of RAxML-NG are provided for MacOS and Linux. A Windows version will be added when available from the RAxML-NG development team (https://github.com/amkozlov/raxml-ng).

RaxmlGUI 2.0 is structured in two parts (Fig. 1), providing on the left all the commands and options to upload input files, set up the analysis, define substitution models and partitions, among other features. On the right panel, it provides options to choose the RAxML version and start the analysis. A RAxML console is integrated in the GUI showing the progress of the analysis, the commands used to launch the analysis and all the screen output produced by RAxML.

**Figure 1:**
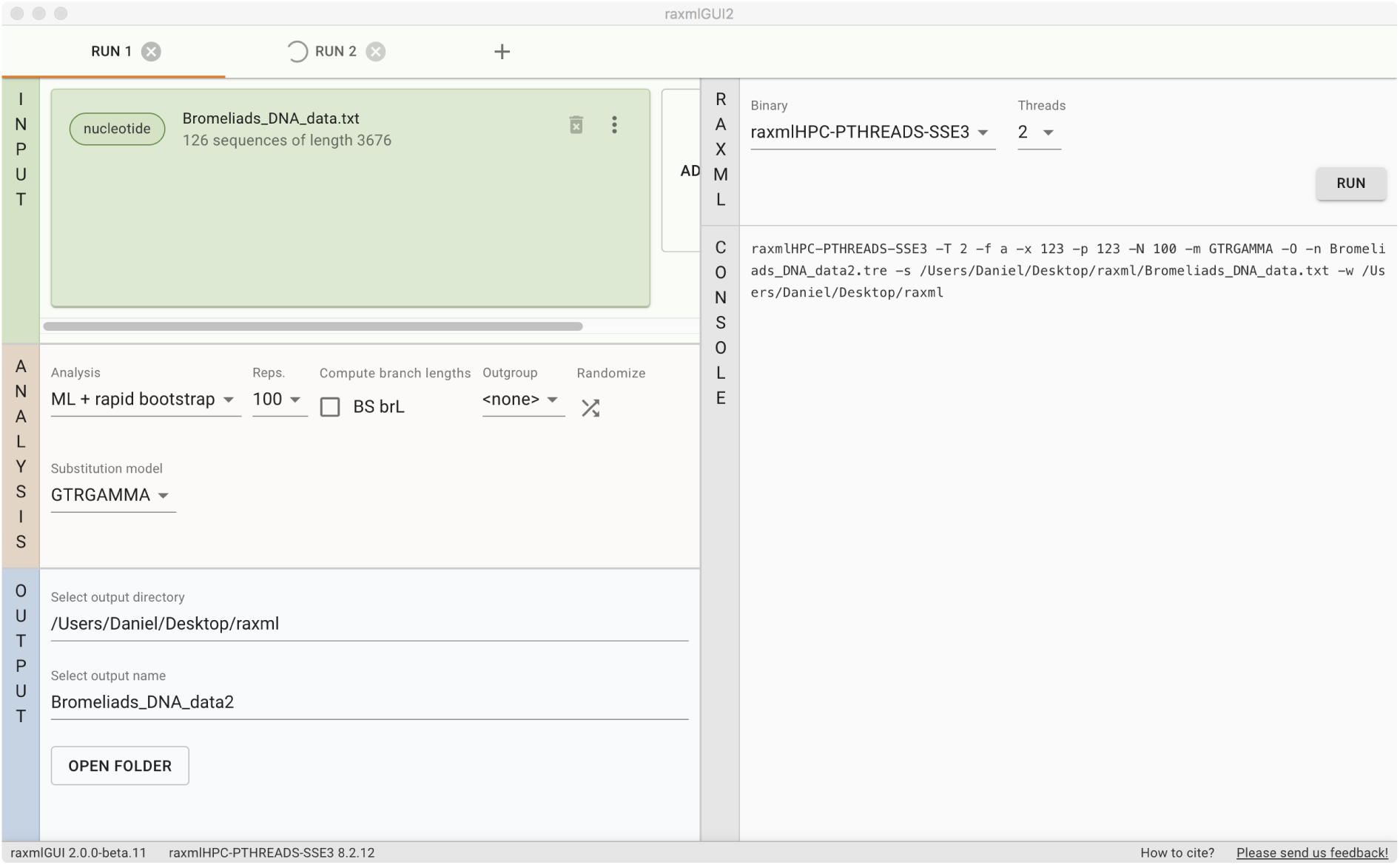
The raxmlGUI 2.0 interface, upon loading an alignment file with amino acid sequences. The input panel provides options to load new alignments and create a concatenated file and to specify partition-specific substitution matrix. The analysis panel provides options to specify the type of analysis, evolutionary models and outgroup selection. The output panel gives easy access to the folder with the input files and a list of output files that appears upon completing the analysis. On the right side of the window the user can select the version of RAxML, start the analysis, and visualize the RAxML output. A tab bar on the top of the window allows users to easily setup multiple runs and switch between them.

### Basic setup

RaxmlGUI 2.0 supports alignment files in two formats: extended PHYLIP and FASTA (example files are available in the program’s repository). Upon loading an alignment, the program parses the names attributed to each sequence (e.g. the species name) and creates a list of taxa in the *Outgroup* menu button, which can be used to root the tree based on a user-defined outgroup (note that maximum likelihood trees can always be re-rooted after the analysis using tree-viewing software such as FigTree (Rambaut, 2012)).

Phylogenetic analyses can be run based on different types of data: nucleotide sequences (DNA, RNA), amino acid sequences, discrete binary and multi-state characters (e.g. used for descriptions of morphological data). Since each data type requires a specific class of substitution models, raxmlGUI automatically recognizes the data type from the loaded input file and provides the user with a drop-down menu showing all the substitution models compatible with the alignment.

### Analytical pipelines

The default analysis includes a maximum likelihood search of the best tree, followed by a bootstrap analysis (option *ML + rapid bootstrap*). Bootstrap support values are then drawn onto the maximum likelihood tree. After loading the alignment file and setting up the preferred substitution model, launching the default analysis only requires hitting the *Run* button on the right panel. Other options are available on the analysis panel to set the number of bootstrap pseudo-replicates. The analysis progress can be monitored on the console panel of raxmlGUI 2.0. When the analysis is completed a list of output files will be available on the output panel. Clicking on the file names will open the files in the user’s default program (e.g. FigTree for tree files). The most important output of this analysis is named “RAxML_bipartitions.*input*.tre” (where *input* is by default the file name of the alignment) and includes the maximum likelihood tree topology and branch lengths with labels reporting the bootstrap scores for each node (bipartition) in the tree. All output files are by default saved in the same directory of the input file.

Several other types of analysis are available in raxmlGUI 2.0. Some analyses integrate multiple calls to RAxML to simplify the user experience in a single pipeline. For instance, the *ML + thorough bootstrap* option launches, in a simple click, a sequence of three RAxML calls to 1) infer the maximum likelihood tree through a user-defined number of independent searches; 2) run a user-defined number of thorough non-parametric bootstrap replicates; and 3) draw the bootstrap support values onto the maximum likelihood tree.

### Automatic concatenation of alignments and partitions

An important feature of raxmlGUI 2.0 is the automated concatenation and partitioning of alignments, which simplifies the analysis of multiple genes or combination of different data types, e.g. amino acids sequences and morphological data. After loading the first alignment, the user can add new ones to concatenate them into a single analysis. Upon loading additional alignments, raxmlGUI 2.0 performs the following tasks:

- Parse the data to determine the data type (nucleotides, amino acids, multistate)
- Parse the taxa names to make sure the concatenation of sequences occurs across matching taxa even if they are listed in different order among input files
- For any mismatch between taxa of different partitions, give option to automatically create sequences of missing data in the concatenated alignment or drop taxa with missing sequences in any partition.
- Set default partitions for the new alignments and re-compute the concatenated partition

These features facilitate the concatenation of different alignment files, the creation of the partition files, and the generation of sparse matrices resulting from the combination of datasets with different and only partly overlapping taxonomic coverage. These tools also reduce the probability of errors stemming from manually merging sequences by matching taxa names. Additionally, raxmlGUI 2.0 provides an intuitive interface to create partitions within a single alignment file, including the possibility to specify codon based evolutionary models for coding nucleotide sequences (Fig. 2).

**Figure 2:**
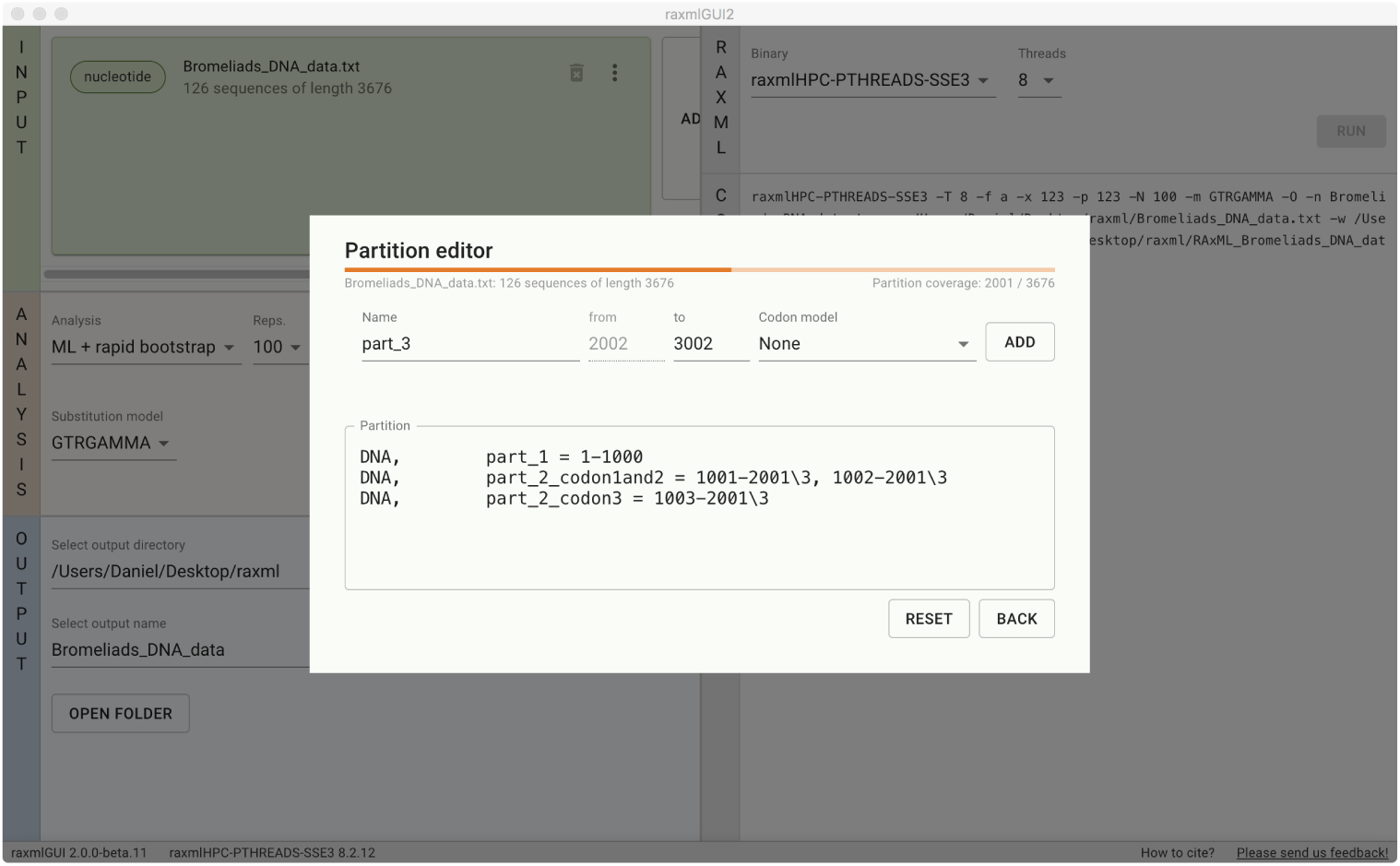
The raxmlGUI 2.0 partition editor. A graphical user interface makes it easy to define a partition for individual alignments.

### Support for both RAxML 8.x and RAxML-NG

In addition to RAxML 8.x, RaxmlGUI 2.0 adds support for RAxML Next Generation (Kozlov et al., 2019), which provides new options and improved performance for very large datasets, which are typical for the analyses of genomic data. Among the novel methods implemented in RAxML-NG, and available through raxmlGUI, is the Transfer Bootstrap Expectation algorithm to quantify topological support for a tree (Lemoine et al., 2018). The user can select which version of RAxML they want to run from the GUI, and the available settings are automatically updated for the specific version. For guidelines of which RAxML version to use for particular objectives and datasets, please refer to (Kozlov et al., 2019).

### Performance and implementation

There is no performance difference between running RAxML on the command line and running it from the GUI as raxmlGUI just forwards all settings as parameters to the command line version of RAxML and runs that as a separate process. RaxmlGUI 2.0 also supports a tabbed interface for running multiple analyses in parallel (Fig. 1).

RaxmlGUI 2.0 is built with Electron (Github Inc, 2020), a framework for creating cross-platform desktop applications using web technologies such as JavaScript, HTML, and CSS. The user interface is built with Material-UI (Material-UI, 2020), a React (Facebook, Inc., 2020) user interface framework with components that implement Google’s Material Design (Google, 2020). The Electron base improves the portability and compatibility across platforms and operating systems compared to the previous version of raxmlGUI that uses an obsolete Python 2.x codebase. The installation is extremely simple and does not require any additional external libraries or dependencies, nor does it require admin rights on the machine.

On machines featuring multiple CPUs (i.e. most desktop and laptop computers) the GUI allows users to easily use RAxML’s powerful parallel computing, which can drastically speed up the analyses. RaxmlGUI 2.0 includes pre-compiled versions of the PTHREAD version of RAxML and a dropdown menu button to specify the desired number of CPUs allocated for the analysis.

### Availability and users’ feedback

RaxmlGUI 2.0 is available at antonellilab.github.io/raxmlGUI/, with links to example files for testing. The program is open source and licensed under a GNU Affero General Public License v3 (AGPL-3.0). We encourage users to report any issues, feature requests, and general feedback either as GitHub issues at github.com/AntonelliLab/raxmlGUI/issues (this requires a GitHub account) or by email to.

## Conclusion

We presented a graphical interface providing an intuitive and user-friendly access to the high-performance phylogenetic software RAxML and RAxML-NG, without compromising performance. Our implementation allows students, professionals, and researchers to use the latest, state-of-the-art methods to build robust phylogenetic hypotheses, irrespective of their computing skills. We hope research and teaching in different fields involving phylogenetic inference -from evolutionary biology to taxonomy, from drug discovery to epidemiology-can benefit from using our program.

## Acknowledgements

We thank I. Michalak and many raxmlGUI users for feedback on the program. D.E. was supported by the Swedish Research Council, Grant No. 2016-00796. D.S. received funding from the Swiss National Science Foundation (PCEFP3_187012; FN-1749) and from the Swedish Research Council (VR: 2019-04739). A.A. is supported by funding from the Swedish Research Council, the Knut and Alice Wallenberg Foundation, the Swedish Foundation for Strategic Research and the Royal Botanic Gardens, Kew.

## Authors’ contributions

DS conceived the ideas; DE, JK and DS designed the interface; DE and JK implemented the ideas and design; DE, JK and DS led the writing of the manuscript. DE, JK, DS and AA contributed critically to the drafts and gave final approval for publication.

